# Spinal motoneurones are intrinsically more responsive in the adult G93A SOD1 mouse model of Amyotrophic Lateral Sclerosis

**DOI:** 10.1101/2020.05.15.098723

**Authors:** D.B. Jensen, M. Kadlecova, I. Allodi, C.F. Meehan

## Abstract

*In vitro* studies from transgenic Amyotrophic Lateral Sclerosis models have suggested an increased excitability of spinal motoneurones. However, *in vivo* intracellular recordings from adult ALS mice models have produced conflicting findings. Previous publications using barbiturate anaesthetised G93A SOD1 mice suggested that some motoneurones are hypo-excitable, defined by deficits in repetitive firing. Our own previous recordings in G127X SOD1 mice using different anaesthesia, however, showed no repetitive firing deficits, and increased persistent inward currents at symptom onset. These discrepancies may be due to differences between models, symptomatic stage, anaesthesia or technical differences. To investigate this, we repeated our original experiments, but in adult male G93A mice at both presymptomatic and symptomatic stages, under barbiturate anaesthesia.

*In vivo* intracellular recordings from antidromically identified spinal motoneurones revealed no significant differences in the ability to fire repetitively in the G93A SOD1 mice. Motoneurones in G93A SOD1 mice fired significantly more spontaneous action potentials. Rheobase was significantly lower and the input resistance and input-output gain were significantly higher in both presymptomatic and symptomatic G93A SOD1 mice. This was despite a significant increase in the duration of the post-spike after-hyperpolarisation (AHP) in both presymptomatic and symptomatic G93A SOD1 mice. Finally, evidence of increased activation of persistent inward currents was seen in both presymptomatic and symptomatic G93A SOD1 mice. Our results do not confirm previous reports of hypo-excitability of spinal motoneurones in the G93A SOD1 mouse and demonstrate that the motoneurones do in fact show an increased response to inputs.

**Key Point Summary:**

- Although *in vitro* recordings using neonatal preparations from mouse models of Amyotrophic Lateral Sclerosis (ALS) suggest increased motoneurone excitability, *in vivo* recordings in adult ALS mouse models have been conflicting.
- In adult G93A SOD1 models, spinal motoneurones have previously been shown to have deficits in repetitive firing, in contrast to the G127X SOD1 mouse model.
- Our *in vivo* intracellular recordings in barbiturate-anaesthetised adult male G93A SOD1 mice reveal no deficits in repetitive firing either prior to or after symptom onset.
- We show that deficits in repetitive firing ability can be a consequence of experimental protocol and should not be used alone to classify otherwise normal motoneurones as hypo-excitable.
- Motoneurones in the G93A SOD1 mice showed an increased response to inputs, with lower rheobase, higher input-output gains and increased activation of persistent inward currents.

## Introduction

Amyotrophic Lateral Sclerosis (ALS) is a neurodegenerative disease, characterised by a progressive loss of upper and lower motoneurones leading to muscle weakness, paralysis and eventually death. However, patients also develop gain of function motor symptoms including fasciculations and spasticity, which suggests an increased excitability of spinal motoneurones. The main drug to have any effect on the survival of patients, Riluzole (Bensimon *et al*., 1994; Miller *et al*., 2012), was initially thought to mediate this effect by blocking glutamatergic neurotransmission (Benavides *et al*., 1985; Centonze *et al*., 1998). In addition to this, it has now also been shown to affect many ion channels including Ca^2+^ -activated BK channels (Wu & Li, 1999; Wang *et al*., 2008), channels mediating both persistent (Hebert *et al*., 1994; Schuster *et al*., 2012) and inactivated transient Na^+^ channels (Urbani & Belluzzi, 2000; Wang *et al*., 2008) as well as voltage gated Ca^2+^ channels including those mediating persistent inward currents (PICs) (Lamanauskas & Nistri, 2008; Bellingham, 2013). This further implicates neuronal excitability in the disease pathogenesis, either by an increased excitation of the spinal motoneurone or by an increased intrinsic excitability of the motoneurone itself.

To test this hypothesis, it is necessary to record directly from the motoneurones. Such experiments were initially performed in *in vitro* preparations from genetically modified mouse models of ALS, generally confirming an increased excitability of motoneurones and showing an increase in PICs (Kuo *et al*., 2004; Kuo *et al*., 2005; Quinlan *et al*., 2011). However, these recordings were generally restricted to embryonic and neonatal tissue. Studies examining intrinsic motoneurone excitability *in vivo* in adult ALS mice have shown conflicting results. Initial investigations in the G127X SOD1 mouse model suggested that increased Ca^2+^ PICs persist into adulthood (Meehan *et al*., 2010a). Additionally, at symptom onset, the axon initial segment elongates and Na^+^ currents appear to be increased in this region (Jørgensen *et al*., Under review). Conversely, studies using the G93A SOD1 and FUS models have reported a lack of repetitive firing in response to a triangular current ramp injection in the soma of some motoneurones which has been interpreted as a hypo-excitability of the motoneurones (Delestree *et al*., 2014; Martinez-Silva *et al*., 2018). Given the recent focus on the development of new treatments influencing neuronal excitability for ALS, such as Retigabine (Noto *et al*., 2016), it is crucial to solve these inconsistencies.

One explanation for the contradicting findings could be in the use of different models. It is always possible that the actual disease mechanisms may be different between the mouse models. Additionally, the timing of the experiments relative to the disease progression may also affect the results. The G127X SOD1 model has a slow onset, but a rapid progression (Jonsson *et al*., 2004), while the G93A has an earlier onset and a slower progression (Gurney *et al*., 1994). Even though both mouse models have been studied at presymptomatic time points, it is likely that experiments were performed at different pre-clinical stages of the disease. This is highly relevant as excitability changes are observed with disease progression in the G127X SOD1 mouse (Jørgensen *et al*., Under review).

Differences in the anaesthetics used in the different experiments may also have contributed to the conflicting results. The experiments with the G93A SOD1 and FUS models used barbiturate anaesthetics (Delestree *et al*., 2014; Martinez-Silva *et al*., 2018) which are known to block PICs (Guertin & Hounsgaard, 1999; Button *et al*., 2006). This may be important if the increase in PICs is a compensatory mechanism to maintain the intrinsic motoneurone excitability in these mice. Consequently, this anaesthetic may preferentially impair firing ability in ALS models.

Finally, as all the *in vivo* studies observing increased excitability in the adult mouse have been produced in one laboratory, while all the *in vivo* studies finding a lack of repetitive firing come from another, technical differences between the way experiments are performed in the different laboratories might explain the differing results. The aim of this study was therefore to repeat our previous studies that we performed in the G127X SOD1 mouse (Jørgensen *et al*., Under review), but this time using the same SOD1 model and anaesthetic as Delestree *et al*. (2014) and Martinez-Silva *et al*. (2018). To control for potential excitability changes with disease stage, experiments were perfomed at both presymptomatic and symptomatic time points. Preliminary data has been published in abstract form (Jensen & Meehan, 2018).

## Methods

All procedures were conducted in accordance with the EU Directive 20110/63/EU and the Danish Animal Inspectorate (Permission no: 2018-15-0201-01426), along with institutional approval from the Department of Experimental Medicine at Copenhagen University and comply with the ethical policies of the Journal of Physiology. SOD1 G93A mice (B6.Cg-Tg(SOD1-G93A)1Gur/J, stock no: 004435, The Jackson Laboratory) were bred at our own institution using male SOD1 G93A and female C57BL6/J mice from Jackson. Mice were fed ad libitum, with constant access to water with a 12:12 hour light/dark cycle. Only male mice were used to eliminate variability in disease onset and progression (Choi *et al*., 2008; Alves *et al*., 2011). All mice were genotyped using DNA extracted from ear clippings and with PCR using the following primers: 5’-CATCAGCCCTAATCCATCTGA-3’ and 5’-CGCGACTAACAATCAAAGTGA-3’ as suggested by The Jackson Laboratory. Furthermore, we tested the transgene copy number using real time qPCR using the following primers: 5’-GGGAAGCTGTTGTCCCAAG-3’ and 5’-CAAGGGGAGGTAAAAGAGAGC-3’ for the SOD1 gene and 5’-CACGTGGGCTCCAGCATT-3’ and 5’-TCACCAGTCATTTCTGCCTTTG-3’ for the reference gene. The primers were mixed with 10ng of extracted DNA and KAPA SYBR® FAST master mix (Merck KGaA, Damstadt, Germany). The qPCR reaction was performed (BIORAD CFX96) with initial temperature of 95°C for 10 minutes followed by 40 cycles of 10 seconds at 95°C and 45 seconds at 60°C. The ΔCT value between the two genes from each G93A SOD1 mouse were compared with ΔCT values from the mice obtained from The Jackson Laboratory and all were within one third of these. Experimental blinding was applied when possible, given the presence of obvious symptoms.

### Behavioural characterisation

To characterise our specific colony of this strain, we first performed a simple cage grip test in 7 male G93A mice and 6 Wild-Type (WT) male littermates (from P63). In this test, the mouse was placed on a wired cage lid which was gently inverted. The endurance time during which they could hang on to the lid before falling to the padded surface below was recorded (tested to a maximum of 2 minutes). The test was repeated 3 times with 10 minutes breaks in between and the mean time was calculated. These mice were followed to humane endpoint (paralysis in one or both limbs). This data was then used to select the age groups for the electrophysiological experiments which were conducted on 18 different male mice: 6 symptomatic mice (mean: 113 days, SD: 3.2, actual range: 107-116), 6 presymptomatic mice (mean: 72 days, SD: 2.2, actual range: 69-75) and 6 WT littermate mice (mean: 73 days, SD: 2.9, actual range: 70-78). The cage grip test was then also performed on these G93A SOD1 mice used for the electrophysiological experiments to confirm their symptomatic stage.

All electrophysiological experiments were terminal and conducted under anaesthesia ensuring no additional load on the animals’ welfare. Mice were painlessly euthanised at the end of the experiments by an overdose of the barbiturate anaesthesia (> 4× the induction dose-see below).

### Intracellular recordings

Mice were briefly anaesthetised with Isoflurane (Baxter) to induce a stress-free longer term anaesthesia using sodium pentobarbital (75 mg/kg I.P.) with supplementary doses given as needed (by testing reflexes to noxious pinch of the hind paw). At the start of the surgery the mice received a subcutaneous injection with atropine (0.02 mg) to decrease mucous secretions. Two catheters were inserted I.P. for the further administration of anaesthetics and the neuromuscular blocker Pavulon (Pancuronium Bromide, 2 mg/ml, diluted 1:10 with saline; an initial dose of 0.1 mL dose followed by a 0.05 mL dose every hour) during the recording procedure (see below). A tracheotomy was performed and a tracheal cannula was inserted for subsequent ventilation. The sciatic nerve of the left hind limb was dissected but left connected to muscles and a hemi-laminectomy was performed at vertebral levels T13 and L1. The mouse was then placed in a stereotactic frame with vertebrae clamps on the T12 and L2 vertebrae. Oil pools were made around the left hind limb and around the spinal cord by retracting the skin with silk threads. The temperature was controlled using a rectal thermometer with feedback to a heating lamp above and a heating pad below the mouse and kept at 37°C. The mouse was artificially ventilated with a mix of oxygen and air at 70 breaths per minute and the expired CO_2_ level was measured using a Capstar CO_2_ analyser (IITC Life Science Inc., Woodland Hills, CA, USA). The electrocardiogram was measured throughout using clips on the ear and the hind paw and was used to monitor the anaesthetic under neuromuscular blockade. The sciatic nerve was placed on bipolar stimulating hook electrodes and a ball electrode was placed on the dorsal side of the spinal cord to record the cord dorsum potential (CDP) resulting from stimulation of the sciatic nerve (0.1 ms pulse, 5 × threshold for the CDP). Intracellular recordings were obtained with sharp glass microelectrodes pulled on a P97 Sutter electrode puller and filled with 2M potassium acetate and had a final resistance of ∼25 MΩ. Impaled cells were confirmed to be motoneurones by antidromic identification using stimulation of the sciatic nerve and the incoming volley from the CDP for timing. The membrane potential was recorded and amplified with an Axoclamp 2A amplifier (Axon Instruments, Union City, CA, USA) and further amplified and filtered using Neurology amplifiers (Digitimer, UK). Signals were then digitized using a CED 1401 digital to analog converter (RRID:SCR_017282, Cambridge Electronic Design, Cambridge, UK) and recorded using Spike2 software (RRID:SCR_000903, Cambridge Electronic Design, Cambridge, UK).

To evoke repetitive firing, triangular current injections were given through the same microelectrode at a speed of 3.3 nA/s using discontinuous current clamp (DCC) mode of the Axoclamp amplifier with switching frequencies of 3 kHz in all motoneurones (in some motoneurones this was also tested at 8 kHz). If the motoneurone did not respond with repetitive firing initially upon penetration, it was tested again later in the recording to allow time for the electrode to seal in the cell. Adequate capacitance compensation of the electrode was manually confirmed immediately prior to all current injections. Only motoneurones with membrane potentials more hyperpolarised than -50 mV at the time measurements were made (confirmed on exit from the cell dorsally) with overshooting antidromic action potentials were included in the analysis.

The analysis was also performed using Spike2 software. In stable recordings, the triangular current injection was used to measure the voltage threshold and rheobase. The input-output measure was measured using an x-y plot of injected current by instantaneous firing frequency (measured in Spike 2). The elevation of the slope (the I-f gain) was calculated during the primary range. In some cells, at the end of the primary range, an inflection was seen representing an increase in gain which is thought to be due to the somatically applied currents now reaching dendritic L-type calcium channels mediating PICs. In those cells, the firing frequency at the onset of the secondary range was measured. Another indirect measure of PICs, ΔI, was also measured as the difference in the current at recruitment on the ascending part of the current ramp and the current at de-recruitment on the descending part of the current ramp.

In some cells, the amplitude and duration of the post-spike after-hyperpolarization (AHP) was measured using averages of single action potentials generated following a brief intracellular current injection in bridge mode. The duration was measured as the time to return to 2/3 of the amplitude (as in (Meehan *et al*., 2010a)). Finally, the input resistance was tested by injecting a 3nA hyperpolarising, 50 ms square pulse in DCC mode at 5 kHz.

### Statistics

Statistical analyses were performed in GraphPad Prism. D’Agostino & Pearson omnibus normality tests revealed that all data sets tested were not normally distributed and so non-parametric tests were used (Kruskal-Wallis with post hoc Dunn’s multiple comparisons). Medians and inter-quartile ranges (Mdn + QR) are therefore used to describe the data. Chi-square tests were used to analyse contingency data. Details for the specific analyses are provided in tables 1 and 2. The level of significance was set at P ≤ 0.05.

## RESULTS

To characterise the SOD1 G93A strain in our facility, we performed a simple standard grip strength test on a separate cohort of mice followed to our humane endpoint. Results of the test can be seen in figure 1B. Based on this test, we chose two time-points to perform the electrophysiological experiments; one group before symptom onset or significant cell loss (∼70 days) and one group after symptom onset (∼115 days) when significant cell loss has been demonstrated (Fischer *et al*., 2004). We also performed the same test on the mice used for electrophysiological recordings on the day of the experiment. The mean values for these are shown on the same graph (figure 1B) to illustrate the disease stage of the mice used for the electrophysiology experiments relative to the general disease progression for our colony of this strain. The younger mice generally performed within the same range of the other mice from our colony. The older mice performed slightly worse than the average for our colony which we assume to be due to the novelty of the task in these mice. Either way, we can classify the two groups as presymptomatic (PS) and symptomatic (S).

**Figure 1:**
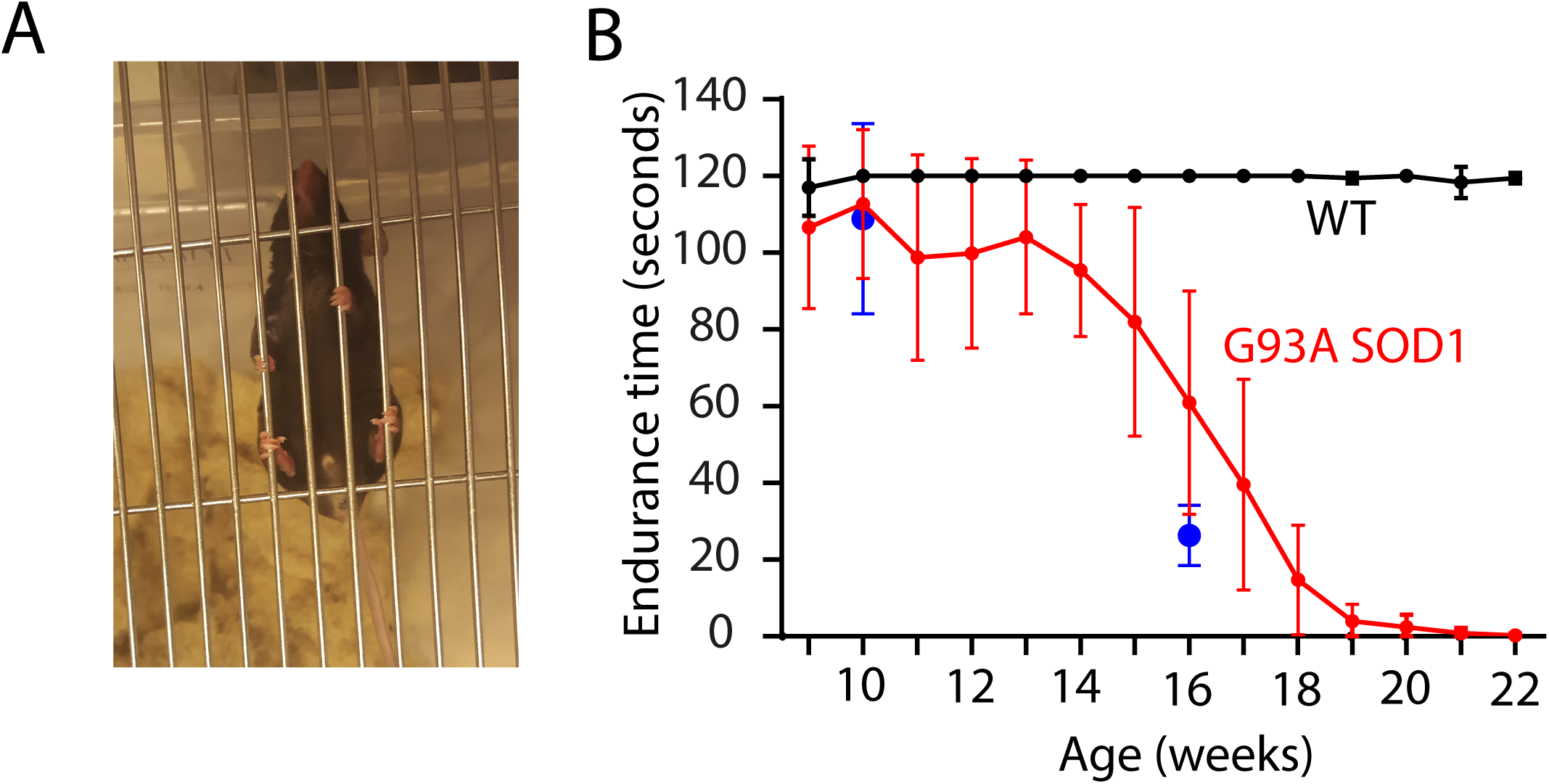
Grip strength test. A. Photomicrograph showing a WT mouse gripping on to an inverted cage lid above a padded surface. The endurance time was measured (up to 2 minutes). B. Graph to show the mean endurance time on the cage grip test for a cohort of 6 WT (black) and 7 G93A mice (red) measured weekly until the G93A mice reached their humane endpoint. This was used for selecting time points for the electrophysiological experiments. Also shown on the graph is the mean for the mice used for the electrophysiological experiments. Error bars show standard deviation (SD).

### Motoneurones show spontaneous activity upon penetration in G93A SOD1 motoneurones

Intracellular recordings were then performed in the mice and penetrations meeting our criteria were made in 329 motoneurones (126 WT, 114 PS, 89 S). In some motoneurones, a spontaneous action potential discharge was observed after entering the cell. An example of this from a PS mouse can be seen in figure 2A. In many cases, it was necessary to inject hyperpolarising current to prevent the firing so that the ramp current injection could be tested. The proportion of cells showing such spontaneous activity was significantly higher in the SOD1 mice (both PS and S) than in the WT mice (WT: 9.5%, PS: 37.7%, S: 49.4%. Chi-square: X^2^ =44.33, p<0.0001, figure 2B).

**Figure 2:**
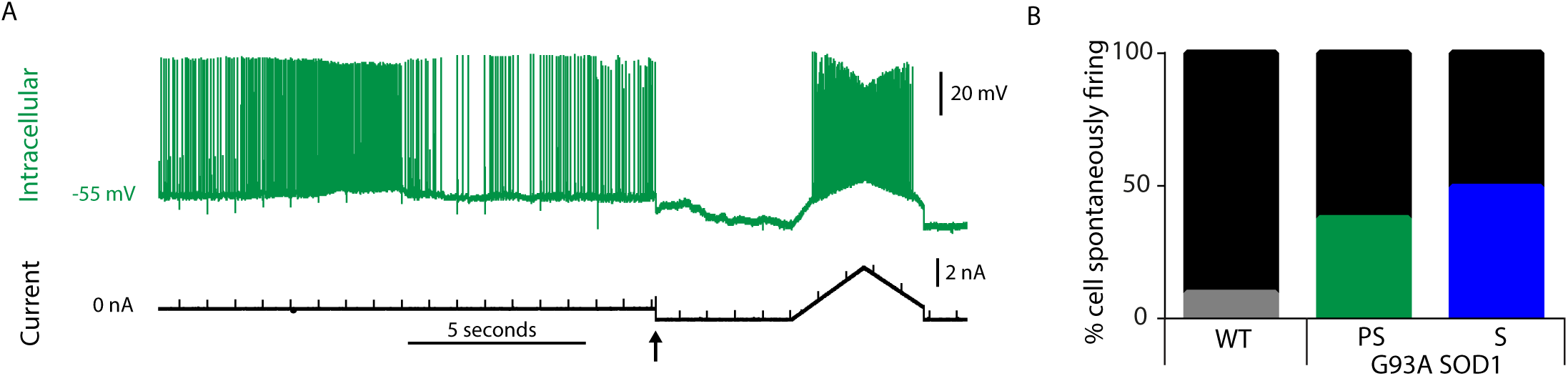
Spontaneous action potential firing in G93A SOD1 mice. A. Example of an intracellular recording (upper trace, green) from a motoneurone in a PS G93A SOD1 mouse soon after initial penetration. The injected current trace (black) is shown below and shows how the spontaneous firing can be terminated with hyperpolarising current (applied from arrow) so that the ramp current injection can be tested. B. The proportion of cells showing spontaneous firing at any point during the recording. This is significantly more frequent in the PS and S G93A SOD1 mice.

### No deficits in repetitive firing are seen in the G93A SOD1 mice

The ability of the motoneurones to fire repetitively in response to an injected triangular current ramp applied to the soma was tested in at least 10 motoneurones per animal and examples are shown in figure 3. In total, 316 motoneurones were tested (119 WT, 109 PS, 88 S) and no obvious deficits in repetitive firing were observed in any of the groups (figure 3). The percentage of cells failing to produce repetitive firing was extremely low and was not significantly different between groups (WT: 3.4%, PS: 0.9%, S: 2.3%. Chi-square: X^2^ =1.571, p=0.4560). However, it is worth noting the major caveat in using a lack of repetitive firing to infer an intrinsic property of a neurone as this property can be easily influenced by technical details. For example, as for the cell illustrated in figure 4A, some motoneurones do not fire repetitively shortly after the initial impalement, but will do so if tested at a later time point once the electrode has had time to seal in. Furthermore, it must be remembered that, *in vivo*, we are using the same electrode to pass current and record using Discontinuous Current Clamp mode on the Axoclamp amplifier. In this mode, the electrode has to quickly switch between passing current and recording which is more difficult to achieve at higher switching frequencies, particularly with the high resistance electrodes used in *in vivo* intracellular recording experiments. Whilst in many cases, repetitive firing is possible to achieve with 8 kHz switching frequency (Figure 4B), despite our best efforts to adequately compensate for capacitance before passing current, occasionally it can be seen that repetitive firing could be obtained in motoneurones using switching frequencies of 3 kHz but not 8 kHz (Figure 4C). This was the case in 8% (17/211) of cells tested at both 3kHz and 8kHz. This was not more frequent in the G93A mice, in fact quite the opposite (WT: 20.6%, PS: 2.8%, S: 1.4% Chi square= 21.36, P<0.0001), suggesting that it was easier to evoke repetitive firing in motoneurones in these mice and so we next turned our attention to determine if this was the case.

**Figure 3.**
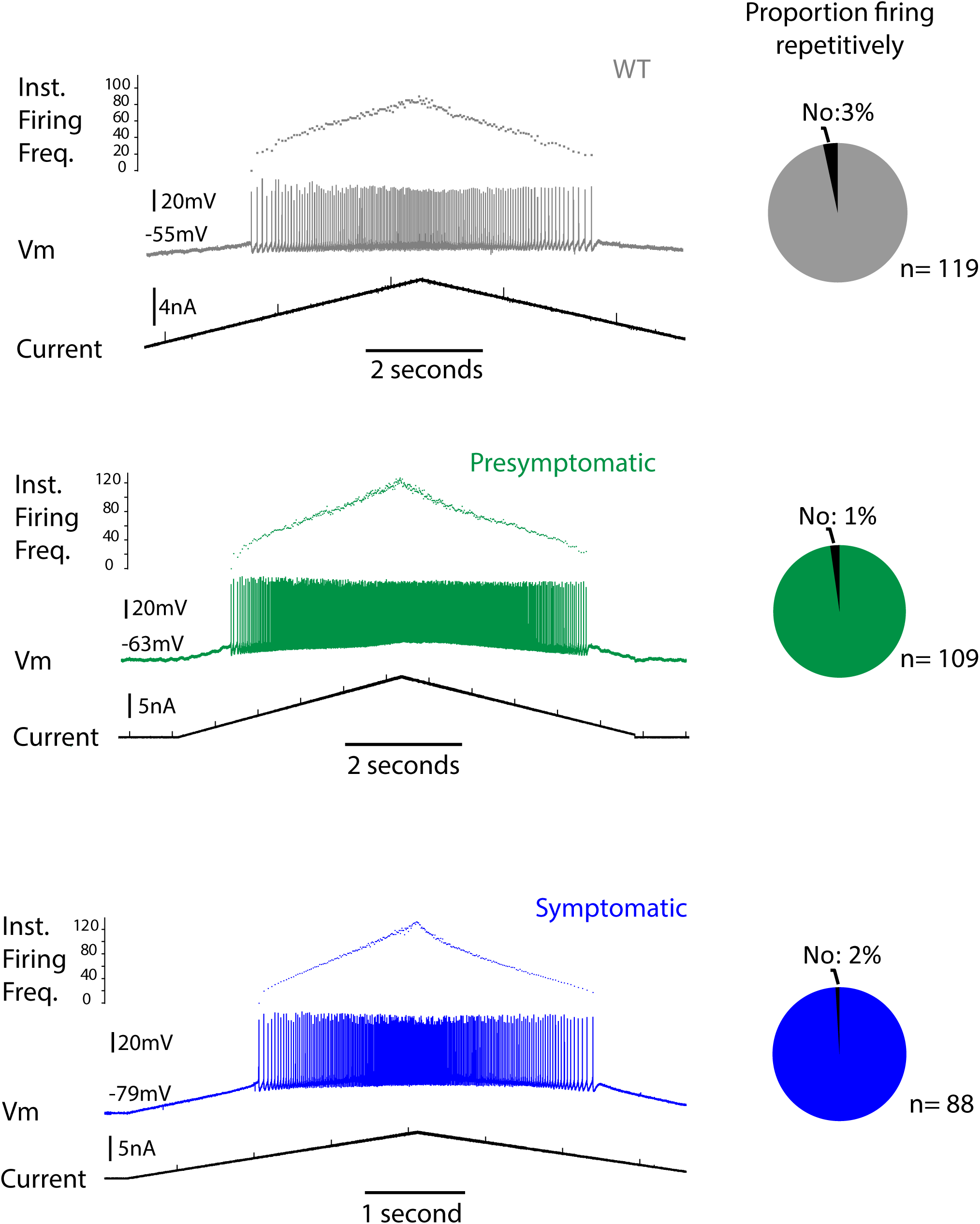
Repetitive firing ability is not impaired in G93A SOD1 mice. Examples of repetitive firing in response to intracellular injection of ramps of current in 3 motoneurones in WT (grey), PS (green) and S (blue) G93A SOD1 mice. To the right of each example is a pie chart showing the proportion of cells in each group responding with repetitive firing which was not significantly different between groups.

**Figure 4:**
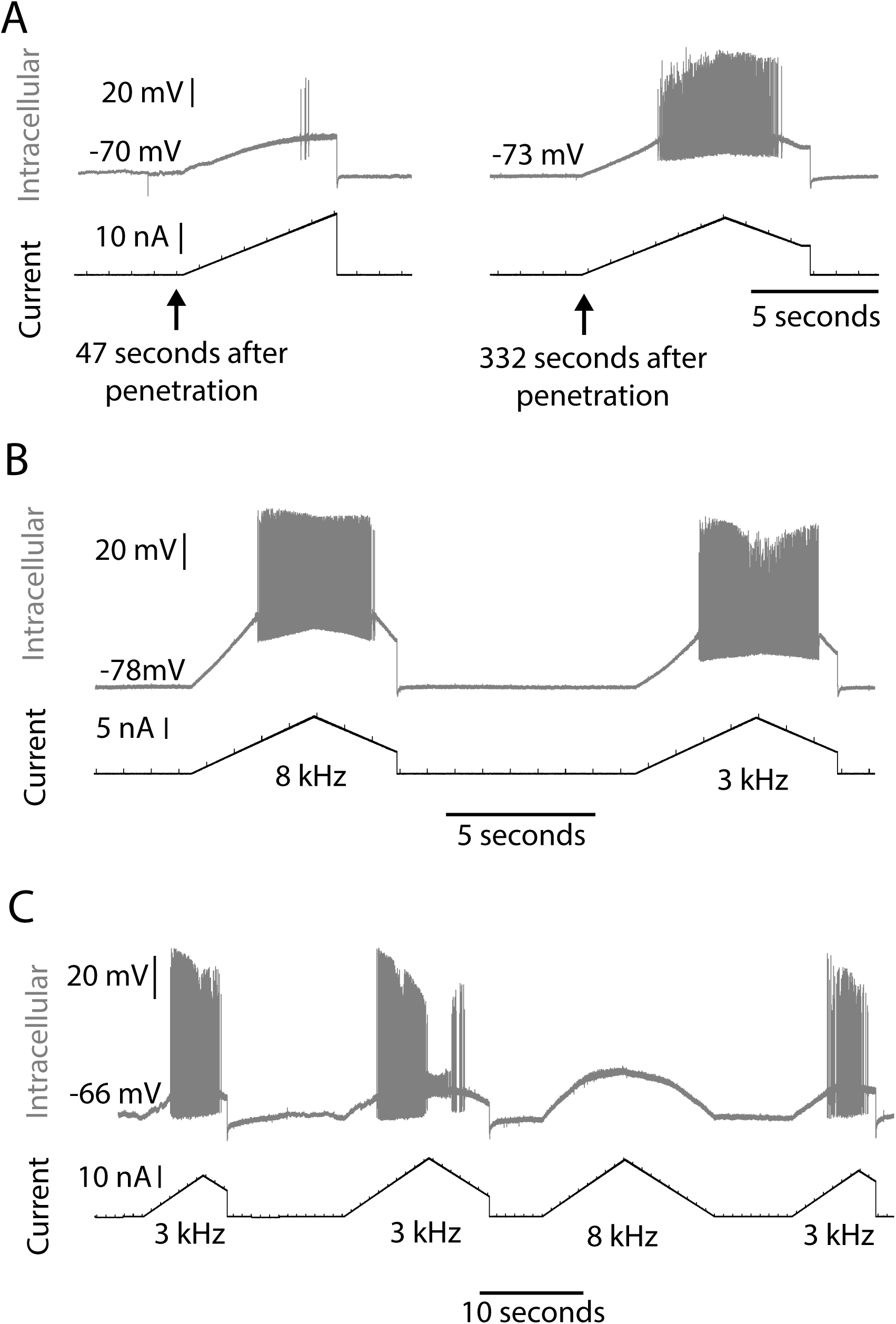
Failure to fire repetitively can be affected by the experimental protocol. A. Example of a recording from a motoneurone that fails to fire repetitively in response to an injection of ramp current 47 seconds after initial penetration. The same neurone, when tested again 285 seconds later, responded with repetitive firing. B. Example of a recording from a motoneurone that fired repetitively in response to an injection of ramp current using both 8 kHz and 3 kHz DCC switching frequency. C. Example of a different motoneurone that fired repetitively in response to an injection of ramp current using 3 kHz but not 8 kHz DCC switching frequency.

### Motoneurones are intrinsically more excitable in the G93A SOD1 mice

The intrinsic excitability of a neurone can be defined by the rheobase current (the amount of current required to elicit an action potential) and, once the neurone is firing, by the input-output gain (the I-f gain). We tested this using the recordings with the injected triangular current ramp. An example of the measurements made is shown in figure 5A. The rheobase in the SOD1 motoneurones was significantly lower in both the presymptomatic and symptomatic group compared to the WT mice (Mdns (+IQR): WT: 6.78nA (6.22), PS: 2.86 nA (2.81), S: 1.80nA (2.96), P<0.001, Figure 5B, Table 1). As can be seen from the spread of individual data points, the overall range is reduced in the G93A mice. To test whether the decrease in rheobase was due to motoneurones having a more depolarized resting membrane potential (Vm) we compared this between groups. The mean Vm was more depolarised in the PS and S mice but this was not significantly different at the P<0.05 level (Mdns (+IQR): WT: -68.94mV (13.78), PS: -64.71mV (11.59), S: -64.98mV (10.7), P=0.0602, Figure 5C, Table 1). Therefore, to test further whether the motoneurones were intrinsically more excitable, we measured the input-output gain as current-firing frequency (I-f) slopes during the primary range of firing. Representative examples of I-f slopes are shown for three motoneurones, one from each group (Figure 5D). The mean I-f gain of motoneurones from the G93A mice was significantly higher than WT for both PS and S mice (Mdns (+IQR): WT: 10.24 Hz/nA (4.79), PS: 12.35 Hz/nA (7.48), S: 15.6 Hz/nA (6.9), P<0.001, Figure 5E, Table 1). As the I-f slope is highly influenced by the AHP, we measured this in 224 cells (WT: 93, PS: 67, S: 64), representative examples of which are shown in Figure 5F. Although the overall range of the amplitude increased, there was only a significant increase in the symptomatic group (Mdns (+IQR): WT: -2.47mV (1.36), PS: 2.87mV (1.72), S: 2.91mV (2.64), P=0.0268, Figure 5G, Table 1), although this could potentially be explained by a more depolarized average Vm. The duration of the AHP was significantly longer in both the PS and S G93A mice compared to the WT and even increased in duration between PS and S mice (Mdns (+IQR): WT: 20.83ms (4.43), PS: 25.92ms (5.64), S: 29.98ms (6.68), P<0.0001, Figure 5H, Table 1). This cannot therefore explain the increase in I-f gain in the motoneurones in the G93A mice and so we next tested the input resistance using the average voltage response to a 3nA, 50ms square current pulse (examples shown in Figure 5I). The input resistance was significantly higher in both PS and S mice than in WT mice (Mdns (+IQR): WT: 3.73 mΩ (3.05), PS: 5.56 mΩ (2.25), S: 6.45 mΩ (3.32) Figure 5J, Table 1).

**Table 1:** Statistical tests for the data shown in figure 5

| Analysis | Cells (n) | % or median (with Quartile values) | Test | Test value | p-value | Post-hoc test | p-value |
| --- | --- | --- | --- | --- | --- | --- | --- |
| Spontaneous firing | WT: 126<br>PS: 114<br>S: 89 | 9.52%<br>37.72%<br>49.44% | Chi-Square | $X^2=44.33$ | <0.0001 | ($X^2=26.93$)<br>($X^2=43.14$)<br>($X^2=2.803$) | WT vs. PS: <0.0001<br>WT vs. S: <0.0001<br>PS vs. S: 0.0941 |
| Repetitive firing | WT: 119<br>PS: 109<br>S: 88 | 96.64%<br>99.08%<br>97.73% | Chi-Square | $X^2=1.571$ | 0.4569 | | |
| Rheobase (nA) | WT: 88<br>PS: 76<br>S: 66 | 6.78 (Q1: 3.59, Q3: 9.81)<br>2.86 (Q1: 1.58, Q3: 4.39)<br>1.80 (Q1: 1.14, Q3: 4.10) | Kruskal-Wallis | K=66.48 | <0.0001 | Dunn's multiple comparisons | WT vs. PS: <0.0001<br>WT vs. S: <0.0001<br>PS vs. S: 0.1469 |
| Resting Vm (mV) | WT: 86<br>PS: 76<br>S: 66 | -68.94 (Q1:-74.01, Q3:-60.23)<br>-64.71 (Q1:-70.39, Q3:-58.80)<br>-64.98 (Q1:-70.26, Q3:-59.56) | Kruskal-Wallis | K=5.619 | 0.0602 |  |  |
| Primary I-F gain (Hz/nA) | WT: 77<br>PS: 74<br>S: 61 | 10.24 (Q1: 7.48, Q3:12.27)<br>12.35 (Q1: 10.30, Q3:17.78)<br>15.6 (Q1: 11.45, Q3: 18.35) | Kruskal-Wallis | H=34.89 | <0.0001 | Dunn's multiple comparisons | WT vs. PS: 0.0001<br>WT vs. S: <0.0001<br>PS vs. S: 0.2390 |
| AHP amplitude (mV) | WT: 93<br>PS: 67<br>S: 64 | 2.47 (Q1: 1.63, Q3: 2.99)<br>2.87 (Q1: 1.83, Q3: 3.55)<br>2.91 (Q1: 1.83, Q3: 4.47) | Kruskal-Wallis | K=7.242 | 0.0268 | Dunn's multiple comparisons | WT vs. PS: 0.1798<br>WT vs. S: 0.0344<br>PS vs. S: >0.9999 |
| AHP 2/3 duration (ms) | WT: 93<br>PS: 67<br>S: 64 | 20.83 (Q1: 19.03, Q3: 23.46)<br>25.92 (Q1: 22.58, Q3: 28.22)<br>29.98 (Q1: 26.50, Q3: 33.18) | Kruskal-Wallis | K=96.08 | <0.0001 | Dunn's multiple comparisons | WT vs. PS: <0.0001<br>WT vs. S: <0.0001<br>PS vs. S: <0.0001 |
| Input resistance | WT: 93<br>PS: 67<br>S: 64 | 3.73 (Q1: 2.46, Q3: 5.51)<br>5.56 (Q1: 4.49, Q3: 6.75)<br>6.45 (Q1: 4.92, Q3: 8.232) | Kruskal-Wallis | K=44.19 | <0.0001 | Dunn's multiple comparisons | WT vs. PS: <0.0001<br>WT vs. S: <0.0001<br>PS vs. S: 0.2706 |

**Figure 5:**
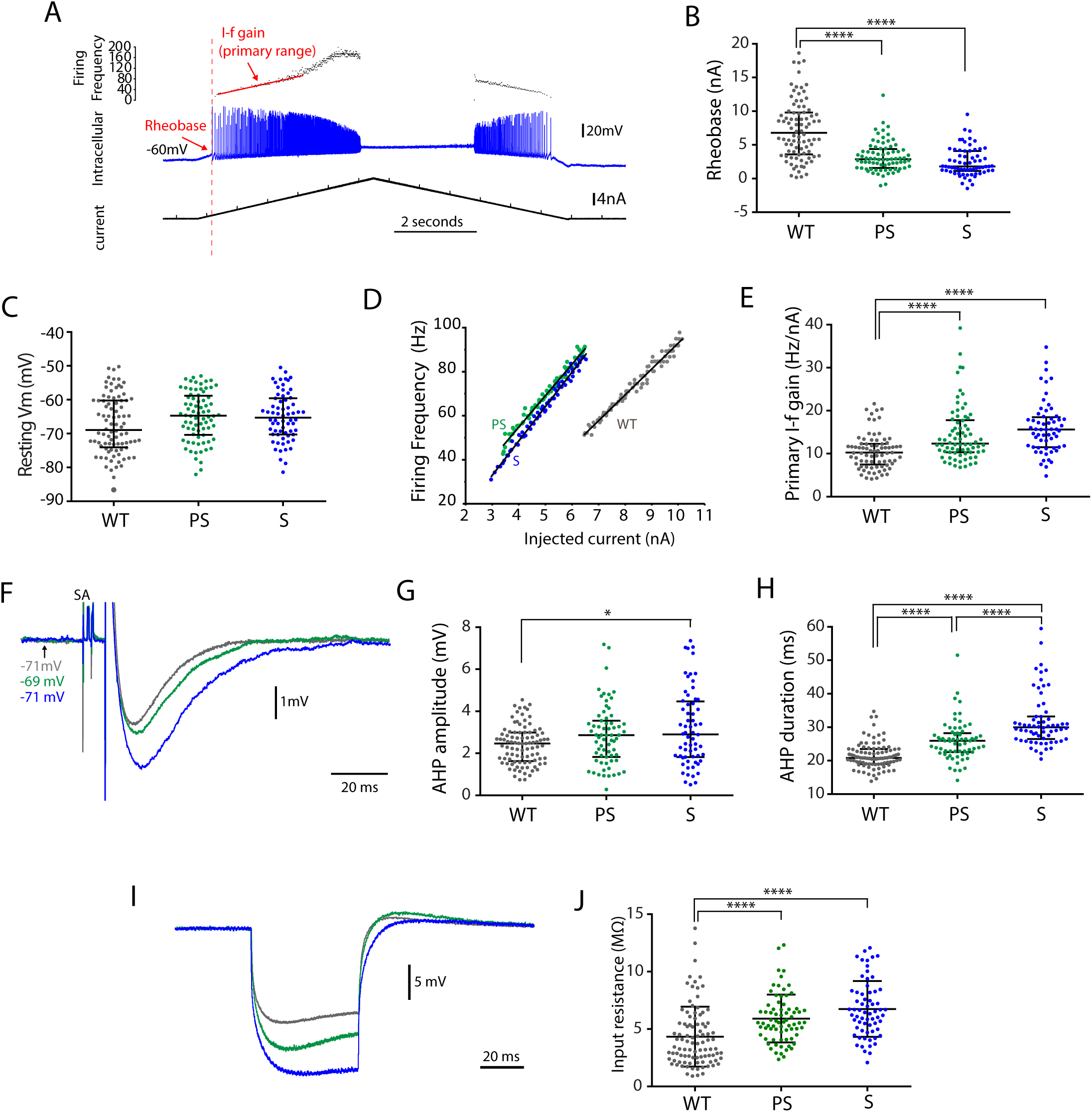
Motoneurones in G93A SOD1 mice are intrinsically more responsive to input. A. Example of an intracellular recording from a motoneurone in an S G93A mouse (blue trace) in response to an intracellular ramp current injection (lower trace) with the instantaneous firing frequency shown in the top trace. Illustrated on the example are the three measurements made including the necessary current for action potential firing (rheobase), the voltage threshold for action potentials and the primary range of firing frequencies used to calculate the I-f gain by plotting the values of firing frequency (top trace) by the current values (bottom trace). B. Scatter dot plot showing individual data points for rheobase (currents necessary to evoke action potential firing) which is significantly lower in both PS and S G93A SOD1 mice. Lines indicate medians and interquartile ranges. C. Scatter dot plot showing individual data points for resting Vm immediately prior to ramp current injection. This is not significantly different between groups. Lines indicate medians and interquartile ranges. D. Representative examples of the current-firing frequency plots (in the primary range) used to measure the I-f gain for three motoneurones from WT (grey), PS (green) and S (blue) mice. E. Scatter dot plot showing individual data points for the I-f gain of motoneurones which was significantly higher than WT for both PS and S mice. Lines indicate medians and interquartile ranges. F. Representative examples of AHPs recorded from motoneurones from WT (grey), PS (green) and S (blue) mice with similar membrane potentials. Traces shown are averages of at least 10 action potentials. G. Scatter dot plot showing individual data points for AHP amplitude for action potentials from WT (grey), PS (green) and S (blue) mice. This was significantly larger in S G93A mice, which also show a much larger range of values. Lines indicate medians and interquartile ranges. H. Scatter dot plot showing individual data points for AHP duration (measured at 2/3 amplitude) for action potentials from WT (grey), PS (green) and S (blue) mice. Lines indicate medians and interquartile ranges. From this, a significant increase in AHP duration can be seen with disease progression. I. Representative examples of voltage drops in response to a 50ms 3nA hyperpolarising current pulse from WT (grey), PS (green) and S (blue) mice. These were used to measure the input resistance of the motoneurones. J. Scatter dot plot showing individual data points for input resistances for motoneurones from WT (grey), PS (green) and S (blue) mice. Lines indicate medians and interquartile ranges. From this, a significant increase in input resistance can be seen in both PS and S mice.

### Evidence for increased persistent inward currents despite barbiturate anaesthesia

During large ramp current injection in spinal motoneurones, an inflection can often be seen on the I-f slope as the secondary range start, indicating the onset of persistent inward currents (figure 6A). This is depressed by barbiturate anaesthesia in mice (Meehan *et al*., 2017) and so was clearly observed in only 16% (n=80 cells) of the WT motoneurones and in these cases was not particularly pronounced. Despite the anaesthetic, secondary ranges were clearly observed in 32% (n=104 cells) of motoneurones from PS mice and 38% (n=97 cells) of motoneurones from S mice, which was significantly different from WT (Chi-square, X^2^ =10.48, P=0.0053). The instantaneous firing frequency at secondary range onset was also significantly lower in motoneurones from both PS and S mice (WT: 130 Hz (14.8), PS: 113 Hz (38), S: 103Hz (28.5), P<0.0005, Figure 6B, Table 2). An additional indirect method to quantify the effects of PICs is to subtract the de-recruitment current from the recruitment current (figure 6C). Caution must be taken with this approach as it is heavily influenced by spike frequency adaptation which is very pronounced in mice motoneurones (Meehan *et al*., 2010b). This value was significantly higher in the motoneurones from both PS and S mice than in WT (WT: -0.85nA (0.99), PS: -0.26nA (1.3), S: 0.08nA (1.05), P<0.0001, Figure 6D, Table 2), again consistent with increased PICs in these mice. Finally, occasionally the effect of the PICs was sufficient to induce self-sustained firing as shown in the example in figure 6E. Here, repetitive firing continues after the end of the current ramp and requires hyperpolarizing current to terminate it. Such examples of self-sustained firing have never been observed in WT mice under barbiturate anaesthesia. These results all suggest an increase in PICs in motoneurones of the G93A mice.

**Table 2:** Statistical tests for figure 6

| Analysis | Cells (n) | % or median (with Quartile values) | Test | Test value | p-value | Post-hoc test | p-value |
| --- | --- | --- | --- | --- | --- | --- | --- |
| Presence of secondary range | WT: 80<br>PS: 104<br>S: 107 | 16.5%<br>31.73%<br>38.144% | Chi-Square | $X^2=10.48$ | 0.0053 | ($X^2= 5.779$)<br>($X^2= 10.37$)<br>($X^2= 0.9096$) | WT vs. PS: 0.0162<br>WT vs. S: 0.0013<br>PS vs. S: 0.3402 |
| Secondary range onset (Hz) | WT: 13<br>PS: 33<br>S: 37 | 130<br>(Q1:119.7, Q3: 134.5)<br>113<br>(Q1:91, Q3:129)<br>103<br>(Q1: 88, Q3: 116.5) | Kruskal-Wallis | H=15.22 | <0.0005 | Dunn's multiple comparisons | WT vs. PS: 0.0246<br>WT vs. S: 0.0003<br>PS vs. S: 0.3131 |
| $\Delta I$ (nA) | WT: 67<br>PS: 71<br>S: 60 | -0.85<br>(Q1: -2.47, Q3: 0.02)<br>-0.26<br>(Q1: -0.99, Q3: 0.31)<br>0.08<br>(Q1: -0.53, Q3: 0.52) | Kruskal-Wallis | H=22.17 | <0.0001 | Dunn's multiple comparisons | WT vs. PS: 0.0106<br>WT vs. S: <0.0001<br>PS vs. S: 0.1798 |

**Figure 6:**
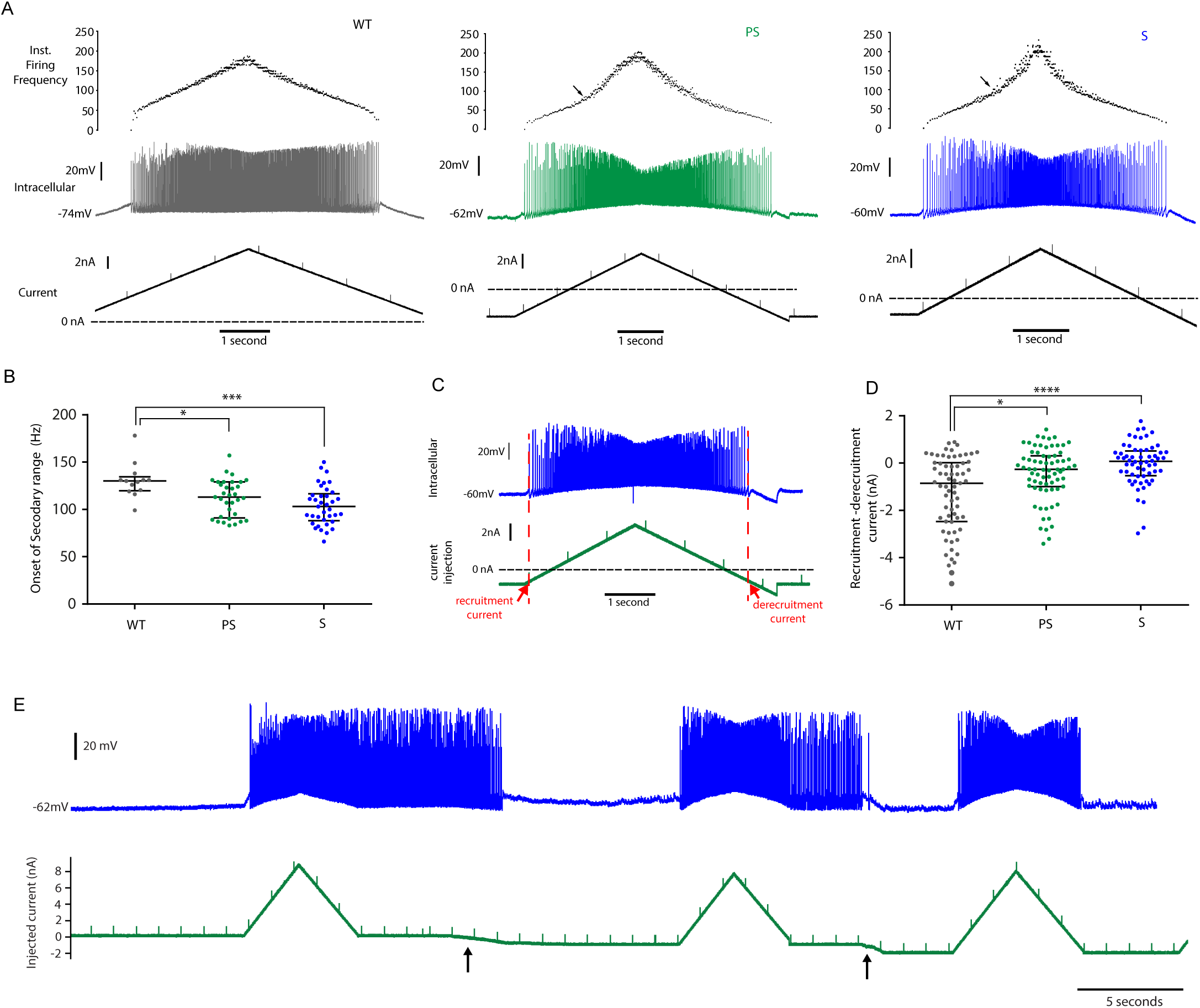
Motoneurones in G93A SOD1 mice show signs of increased activation of persistent inward currents. A. Examples of repetitive firing in response to intracellular injection of ramps of current in 3 motoneurones in WT (grey), PS (green) and S (blue) G93A SOD1 mice. In these examples, the current is taken high enough to reach the secondary range of firing, if present. Here, no sudden increase in gain is visible on the WT recording but clear inflections (arrows) consistent with the onset of the secondary range can be seen in the examples from PS (green) and S (blue) motoneurones. B. Scatter dot plot showing individual data points for the onset of the secondary range for those motoneurones that showed this in WT (grey), PS (green) and S (blue) mice. From this, it can be seen that the onset is at significantly lower firing frequencies in G93A mice. Lines indicate medians and interquartile ranges. C. Example of an intracellular recording from a motoneurone in a S G93A mouse (blue trace) in response to an intracellular ramp current injection (lower trace). In this example, both the current level for recruitment and de-recruitment of the motoneurone is indicated. The first measurement can be subtracted from the second measurement to give an indirect indication of PIC activation. D. Scatter dot plot showing individual data points for recruitment - derecruitment currents for motoneurones from WT (grey), PS (green) and S (blue) mice. Lines indicate medians and interquartile ranges. These values are significantly greater in the G93A mice consistent with increased PIC activation. E. Example of an intracellular recording from a motoneurone in a S G93A mouse showing how the PICs can induce self-sustained firing. Note that the repetitive firing continues after the current ramp is finished. This self-sustained firing can be terminated by applying hyperpolarizing current (at arrows).

## Discussion

The results of these experiments are relatively consistent to those we have previously obtained in the G127X SOD1 mouse using the same protocol but with different anaesthetics. This includes no loss in the ability to fire action potentials repetitively (even in symptomatic mice) and increases in persistent inward currents (Bonnevie *et al*., 2020; Jørgensen *et al*., Under review). We also saw signs of an increased responsiveness to input seen as a decrease in rheobase and an increased I-f gain.

### Repetitive firing and the interpretation of hypo-and hyperexcitability

We did not find evidence of hypo-excitability in these experiments. We define hypo-excitability as a decreased output with respect to input which would be evident as a decrease in rheobase or decreased I-f gain; none of which have been observed either in this study or the previous G93A SOD1 study (Delestree *et al*., 2014; Martinez-Silva *et al*., 2018). In fact, mean I-f gains were also higher by a similar magnitude in the G93A mice in Martinez-Silva *et al* (2018), but the number of cells analysed in each group were significantly lower than ours, reducing statistical power. Therefore, our results are not necessarily conflicting. The main difference then between the studies is in the observed inability of some cells to fire repetitively in response to intracellular current injection. This naturally leads to the question as to how a lack of repetitive firing should be interpreted. If a motor unit cannot fire repetitively then it will be unable to obtain tetanic contractions and should perhaps be considered a dysfunctional motoneurone rather than “hypo-excitable”. However, we would also express caution with this conclusion as we have shown that, with sharp microelectrode recordings, the ability to fire repetitively can be heavily influenced by experimental conditions which can potentially explain inconsistent observations between laboratories.

We show that the timing and nature of the current injection can influence repetitive firing. Failure to initiate repetitive firing immediately after the membrane is ruptured and penetrated by a sharp microelectrode can represent an injury response. It is our experience that, if tested again later after the electrode has had the time to “seal in”, these cells almost always regain repetitive firing ability. Furthermore, the type of electrodes necessary for *in vivo* intracellular recording are of high resistance which, in our experience, not only often struggle to handle high switching frequencies in DCC mode but can easily become partially blocked as they penetrate the membrane and consequently have difficulty passing the current. In such cases, one can exit the cell dorsally, clear the electrode and re-enter the cell with current injection now evoking repetitive firing. If there is difficulty passing current, a slow activation of subthreshold sodium currents can lead to spike inactivation. This may also be influenced by the speed of the current ramp that varies considerably between laboratories and also sometimes within studies (e.g. ranging from 0.1nA/s to 5nA/s in Martinez-Silva *et al*. (2018)). These points are all crucial to consider as they would be more likely to affect the cells with the highest rheobase (i.e. fast motoneurones), making these the most difficult to get to fire repetitively under these conditions, which could potentially explain the higher proportion of fast motoneurones in the “non-firer” category for both WT and ALS mice in other studies (Martinez-Silva *et al*., 2018). We therefore conclude that, with *in vivo* intracellular recordings using sharp microelectrodes, a lack of initial repetitive firing, by itself (especially soon after penetrating the cell) should not be taken to indicate a hypo-excitability of motoneurones especially if other excitability measures indicated by other standard measures (e.g. I-f slope, input resistance and rheobase) appear normal or indicate increased excitability.

That some studies still show an increased tendency for failure of repetitive firing in ALS may simply be the unveiling of an increased vulnerability of the motoneurones in the ALS models to the initial penetration and the recovery time, which would not be surprising. However, it should be noted that in Martinez *et al*. (2018) the proportion of “non-firers” in the control group for the FUS mice was in fact larger than the proportion seen in the G93A ALS mice in the same study. This suggests that we must also exercise caution with this conclusion. However, such a hypothesis could also help to explain findings from *in vitro* studies. It would not be surprising for ALS motoneurones to also be more vulnerable to *in vitro* cell culture conditions. This would help to explain results from experiments using induced pluripotent stem cells (iPSC) derived from ALS patients which have been shown to demonstrate an apparent increased excitability at early stages (Wainger *et al*., 2014), followed by a loss of repetitive firing in older cultures (Sareen *et al*., 2013; Devlin *et al*., 2015). Mitochondrial dysfunction, coupled with a tendency to increased oxidative stress, would undoubtedly lead to an increased vulnerability of the motoneurones to the *in vitro* conditions and it is therefore important to distinguish between a sick and dying motoneurone and what constitutes a true “hypo-excitability” phenotype that would appear under normal *in vivo* conditions with synaptic activation. In this case, we would argue again in favour of the traditional parameters of an increase in rheobase or decreased I-f gain as better indicators.

The confound of an additional vulnerability of the ALS motoneurones to experimental conditions makes *in vivo* experiments all the more crucial in this disease. If anything, under *in vivo* conditions as the disease progresses, we observed an increase in excitability consistent with our recordings in the G127X SOD1 mice. Although direct recording from motoneurones is not possible in humans, our results would definitely be more consistent with the “upper motor signs” of spasticity observed in both ALS patients and ALS mouse models indicating a homeostatic response of the motoneurones to a reduction in the descending motor drive. This hypothesis is supported by our observations in the G127X mouse model of ALS where the axon initial segments of the spinal motoneurones increase in length upon symptom onset (Jørgensen *et al*., Under review), similar to our observations following spinal cord injury (Dimintiyanova *et al*., 2016).

Our observations of an increase in AHP duration with disease progression are also consistent with our previous results in the G127X SOD1 mouse (Maglemose *et al*., 2017) and the increase in AHP duration observed from the most affected muscles in ALS patients (Piotrkiewicz & Hausmanowa-Petrusewicz, 2011). An increased intrinsic motoneurone excitability would make the motoneurones easier to recruit, but would provide less gradual gain control of the individual motor unit if the I-f gain was too steep. We hypothesise that the increased AHP duration seen in ALS represents a homeostatic response to try to normalise the I-f gain of the cell once it is recruited. This could also be effectively achieved in a task dependent manner by plasticity in the C-bouton system, changes to which have already been implicated in ALS (Pullen & Athanasiou, 2009; Herron & Miles, 2012; Casas *et al*., 2013).

One striking feature of our results was the large increase in input resistance in the G93A mice. Although this could be interpreted as we may not recording from fast motoneurones, this explanation seems unlikely given our large sample size and the fact that, with sharp electrodes, intracellular recording is generally biased to larger cells, we believe that this is unlikely to be merely a sampling issue. Of course, in the symptomatic mice, cell death of the higher threshold, low input resistance fast motoneurones could result in a biased sample, however this explanation can be discounted by the decreased rheobase and increased input resistance we also saw in the presymptomatic mice at an age where there is not significant motoneurone loss in this model (Fischer *et al*., 2004). A highly likely explanation for the increased rheobase may be alterations in soma size. Although spinal motoneurones are enlarged in this model at around 30 days of age compared to WT, this pattern is gradually reversed between then and end stage, at which point a 40% reduction in soma size is observed (Dukkipati *et al*., 2018). This would undoubtedly affect the input resistance and excitability as modelling based on these results demonstrated that size changes of even 20% could produce pronounced changes in rheobase.

The observation of temporal changes in soma size is also relevant for comparison of our results with Delestree *et al*. (2018). In those experiments, mice were used ranging from 38–82 days old, collapsed as a single group, which may have masked emerging increased excitability in older mice. Our results do not rule out a possible decreased excitability in younger mice, but we can conclude that immediately prior to and after symptom onset motoneurones in this model are intrinsically more excitable.

### Role of the increased PICs

One noticeable observation was the evidence of increased PICs seen in these motoneurones. This was particularly surprising given that barbiturate anaesthetics are known to block PICs (Guertin & Hounsgaard, 1999; Button *et al*., 2006), which suggests that the actual increase under non-anaesthetised conditions must be even more than we have observed here. When we initially embarked on this study, we had anticipated that increased PICs in ALS motoneurones would be important for the motoneurone to maintain repetitive firing and that we would also therefore see a loss of repetitive firing in some cells due to barbiturate anaesthetics blocking the PICs.

Increased Na^+^ PICs which have been shown in this model (Delestree *et al*., 2014) and presumably arise from the area with the highest density of Nav 1.6 channels, the axon initial segment, which has been shown to increase in length at symptom onset in the G127X model (Jørgensen *et al*., Under review). These would already be partially activated sub-spike threshold and so, as mentioned earlier, would be critical for kicking off repetitive firing. Ca^2+^ PICs, in contrast, are hypothesised to have a functional location on proximal dendrites (Elbasiouny *et al*., 2005; Bui *et al*., 2006), which explains why they are only activated at higher frequencies with artificial current injection directly to the soma producing the secondary range. Normally, their dendritic location means they can be activated earlier by subthreshold EPSP and amplify the input (Lee & Heckman, 1996). Thus, increases in Ca^2+^ PICs would provide an important increased amplification of the reduced descending excitatory motor commands to the motoneurones under normal conditions. Presumably, this amplification is not specific and would also amplify sensory inputs to the motoneurones contributing to development of hyperreflexia in a similar manner to spinal cord injury (Bellardita *et al*., 2017). From this perspective, the increased intrinsic excitability of the motoneurone would simply be a mechanism to maintain a normal level of network excitability, although this would result in the unfortunate side effect of hyperreflexia.

Finally, whether this increase in intrinsic excitability contributes in any way to the ultimate cell death is unknown, although the ability of Riluzole to block both Na^+^ and Ca^2+^ PICs suggests that this is a potentially important hypothesis to test. Importantly, we also show an increased response of the neurone to artificially applied current directly to the soma. To determine the possible role of this in excitotoxicity, it will be crucial to confirm whether the motoneurones genuinely are more active *in vivo* in awake freely moving mice or if the increased intrinsic excitability is precisely tailored to compensate for the loss of descending drive producing an overall normal functional level of motoneurone excitation. In such a scenario, therapies designed to further decrease motoneurone excitability may actually prove detrimental as they would risk reducing residual motor function.

## Author Contributions

***DBJ:*** acquisition, analysis and interpretation of data. Co-wrote manuscript

***MK:*** acquisition, analysis and interpretation of data (behavioural tests)

***IA:*** acquisition of data (established genotyping and copy number verification)

***CFM:*** acquisition and interpretation of data. Co-wrote manuscript

## Acknowledgements

This work was funded by project grants from the Independent Research Fund Denmark (8020-00330A) and Læge Sofus Carl Emil Friis og Hustru Olga Doris Friis Legat. We thank Dr Peter Kirkwood for critical feedback on previous drafts.

